# Recombination detected in the Heat Shock Protein 90 (HSP90) of the *Bemisia tabaci* species complex

**DOI:** 10.1101/655233

**Authors:** Tonny Kinene, Bruno Rossito De Marchi, Titus Alicai, Livingstone S. Luboobi, Christopher Abu Omongo, Anders Savill, Laura M. Boykin

## Abstract

**Background:** *Bemisia tabaci* (whiteflies) are a global insect pest causing billions of dollars in damage each year, leaving farmers with low yields. In East Africa, whiteflies are superabundant and present on cassava plants throughout the year. Whiteflies do not decrease in number in the hot dry seasons in East Africa, therefore, it has been suggested that the synthesis of Heat Shock Protein (HSP) may protect the whitefly from heat stress and other biotic factors. In this study we used data sequence generated from individual whiteflies to assess variability and recombination of the HSP90 gene in members of the *B. tabaci* species complex.

**Results:** A total of 21 samples were sequenced on Illumina Hiseq 2500 and Hiseq 4000. These included eight genetic groups of *B. tabaci*: 7 SSA1, 5 SSA2, 2 Australia I (AUSI), 2 New World Africa (NWAfrica), *B. afer*, Uganda, Mediterranean (MED), and Middle East Asia Minor 1 (MEAM1). An alignment of 21 HSP90 sequences was generated after mapping and *de novo* assembly. Recombination analysis was performed on an alignment of 27 HSP90 sequences (we added an additional 6 sequences from GenBank). There were 18 recombination events detected in the HSP90 gene of the *B. tabaci* species complex, 7 of which were regarded as events that could be caused by evolutionary mechanisms such as gene duplication other than recombination. The phylogenetic analysis carried out on dataset without recombination events revealed a tree pattern with short terminal branches.

**Conclusion:** Recombination events were detected for members of the *B. tabaci* species complex in the HSP90 gene. This could explain the variability in the HSP90 gene of the *B. tabaci* species complex and highlight the phenomenon of the increased chance of survival and reproductive abundance of whiteflies in hot conditions in East Africa, since recombination is a major driving force of evolution.

## BACKGROUND

The *Bemisia tabaci* species complex (whiteflies) is a group of small phloem sap-feeding insects capable of causing extensive crop damage globally [1, 2]. The species complex is composed of at least 34 species that are morphologically indistinguishable [3, 4]. These species cause extensive damage to plants through direct feeding by both juvenile and adult stages of the whitefly [5-7]. They also cause indirect damage through excretion of honeydew, which covers the leaf and fruit surfaces promoting the growth of black sooty mold. This mold interferes with plant photosynthesis, affects plant growth, and leads to poor quality fruits [8]. Whiteflies also transmit plant viruses in the process of feeding from one plant to another [9]. Yield losses due to direct feeding and black sooty mold are estimated at 40% [8]. In Africa, whiteflies are a major threat to food security [10].

Many smallholder farmers in East Africa rely heavily on cassava as a food security and cash crop because of its tolerance to low fertility soils, low rainfall, low labour intensity and long harvesting window [11-13]. Cassava roots are eaten and are rich in carbohydrates, and the leaves are rich in protein and are eaten as vegetables. Cassava can be processed into other commercial products such as flour, cakes and alcohol. Unfortunately, cassava is highly vulnerable to both the whiteflies and the viruses they transmit. Attempts to protect cassava from whiteflies with chemical pesticides have been ineffective and costly [14-16]. The honeydew excreted by superabundant populations of whiteflies in East Africa interferes with photosynthesis and reduces the effectiveness of insecticides [1, 5, 7, 8]. In Africa, whiteflies transmit two cassava viral diseases: cassava brown streak disease (CBSD) and cassava mosaic disease (CMD) [12, 17-20].

Agriculture development in Africa has been constrained by rising temperatures due to global warming [21]. Impacts are economically significant in countries in East Africa that are heavily reliant on natural resources, such as rainwater, for agriculture. Increasing temperature has resulted in prolonged drought seasons, which have left most plants dry. However, the drought tolerant plants like cassava, which save farmers from famine, are now heavily infested with whiteflies. This has caused loss of food and income for farmers, due to the viral diseases associated with whiteflies that infect their crops. The high numbers of whiteflies seen in cassava fields in East Africa, termed as superabundant populations, have increased with increasing temperatures over time which may be due to the fact that temperature determines the geographical distribution and reproductive abundance of species [22-24].

The capacity of Sub-Saharan Africa (SSA) whitefly species to breed in large numbers during periods of high temperatures in East Africa indicates that SSA species might have developed a molecular response to tolerate heat stress. It is known that high temperatures in the environment can cause organisms to respond with physiological, biochemical or behavioural traits that enhance their chances of survival [23, 25]. During stressful conditions the cells of an organism produce a set of conserved proteins called heat shock proteins (HSPs). HSPs are molecular chaperones that bind to and stabilize unfolded proteins and are found in nearly all living organisms [26, 27]. HSPs are named according to their molecular weight (for example, HSP100, 90, 70, 60, and 40) and not all are associated with the heat shock response [26]. HSP70 and HSP90 are the most widely-studied families and are synthesised when the environmental temperature exceeds the optimal temperature of an organism [26, 28].

Gorovits & Czosnek (2017) revealed that HSP70 and HSP90 in plants and vectors are very important for efficient virus infection. HSP90 is also considered to promote virus replication by interacting with virus replicase in *Bambo mosaic* virus [29]. Differential expression studies on HSP on Middle East Asia Minor 1 (MEAM1), one of the whitefly species in the complex, indicated that heat shock proteins (HSP20, 70 & 90) were up-regulated when whiteflies were subjected to a high temperature of 40° C, and this increased the fitness of whiteflies following heat stress [30]. Other studies indicate that up-regulation of HSP could be key in the determination of the natural geographical distribution of natural populations by seasonal dynamics [31]. HSP response was examined on silverleaf whitefly and it was found that HSP70 and HSP90 were the major polypeptides synthesized by whiteflies in response to heat stress [32]. There is more evidence that HSPs are up-regulated by stress conditions in *B. tabaci* [33-35]. However, none of these studies have been carried out on SSA whitefly species.

In this study we investigated the variability of the HSP90 gene in the *B. tabaci* species complex and found evidence of recombination in the coding region of HSP90 gene in the *B. tabaci* species complex.

## RESULTS

A total of 21 samples were sequenced, which included eight genetic groups of *B. tabaci*: 7 SSA1, 5 SSA2, 2 Australia I (AUSI), 2 New World Africa (NWAfrica), *B. afer*, Uganda, Mediterranean (MED), and Middle East Asia Minor 1 (MEAM1). 15 of these samples were sequenced on an Illumina Hiseq 2500 on a rapid run mode, resulting in 8,000,000 raw reads, and 6 samples were sequenced on an Illumina Hiseq 4000 on a high throughput mode which yielded raw reads ranging from 43,194,264 to 37, 306,246. After trimming for quality, the reads reduced to a range of 7,876,283 to 7,816,022 for the first 15 samples and 42,953,727 to 37,071,584 for the 6 samples (additional file 2). *De novo* assembly produced contigs ranging from 2530 – 30716, which were then mapped to the HSP 90 EU934241 reference sequence. The final coding sequence of length of 2160 base pairs was obtained and used in this study as the HSP90 sequence. The final HSP90 sequence consisted of a consensus between the *de novo* contig of interest and the mapped consensus.

18 recombinant events were identified by RDP4 [36] and 7 of these were flagged as events caused by evolutionary processes such as gene duplication other than recombination (Table 1). These included: SSA2_191_HSP90, SSA2_193_HSP90, AUSI_347_1_HSP90, AUSI_347_2_HSP90, NWI_156_2, NWI_156_3_HSP90 and are shown in (Figure 1).

**Table 1.** A summary of results of recombination analysis from various recombination-detection tests, using RDP4 software

| Event | Recombinant Sequence(s) | Major parent | Minor parent | Detected method in RDP4 | Detected Break-point Positions | Highest P-value |
| --- | --- | --- | --- | --- | --- | --- |
| 1 | SSA1 65 HSP90 | NW1 156 3 HSP90 | SSA2 179 HSP90 | R G B M C S T | 569 - 1682 | 1.465x10<sup>-25</sup> |
| 2 | ^SSA2 191 HSP90 | Unknown | SSA1 189 HSP90 | R G M C S T | 1101 - 1762 | 8.489x10<sup>-20</sup> |
| 3 | ^NW1 156 3 HSP90 | Unknown | SSA1 189 HSP90 | R G B M C S T | 132* - 542 | 4.575x10<sup>-14</sup> |
| 4 | SSA1 70 HSP90 | SSA1 189 HSP90 | Unknown | R G B M C S T | 1927 - 2077* | 1.134x10<sup>-13</sup> |
| 5 | MED 320 3 HSP90 | SSA2 193 HSP90 | SSA1 68 HSP90 | R G M C S T | 1940 - 2160* | 4.243x10<sup>-13</sup> |
| 6 | ^SSA2_191_HSP90 | NW1_156_3_HSP90 | Unknown<br>(ASIAII_3_HSP90_HM013713) | RG_BMCST | 674 - 856 | 2.60x10<sup>-11</sup> |
| 7 | AUS1 347 1 HSP90 | MEAM1 153 HSP90 | NW1 156 3 HSP90 | R G B M C S T | 1884 - 2074* | 4.730x10<sup>-11</sup> |
| 8 | MEAM1 153 HSP90 | MED HSP90 HM013710 | MW1 156 3 HSP90 | R G B M C S T | 574* - 1030 | 5.032x10<sup>-10</sup> |
| 9 | AUS1 347 1 HSP90 | AUS1 347 2 HSP90 | NW1 156 3 HSP90 | G B M C S T | 896 - 1036 | 9.11x10<sup>-10</sup> |
| 10 | MED 320 3 HSP90 | NW1 156 3 HSP90 | MEAM1 HSP90 EU934241 | G B M C S T | 1091 - 1752 | 9.735x10<sup>-10</sup> |
| 11 | SSA2 191 HSP90 | SSA2 193 HSP90 | SSA2 179 HSP90 | R G M C T | 2057 - 2160* | 2.50x10<sup>-8</sup> |
| 12 | ^SSA2 193 HSP90 | MED 320 3 HSP90 | AUS1 347 2 HSP90 | R G B M C T | 318* - 504 | 2.95x10<sup>-8</sup> |
| 13 | ^NW1 156 2 HSP90 | NM1 156 3 HSP90 | ASIAII 3 HSP90 HM013713 | R G M C T | 1076 - 1257 | 4.43 x10<sup>-8</sup> |
| 14 | NW1 156 2 HSP90 | NW1 156 3 HSP90 | MEAM1 HSP90 DQ093380 | G B M C S T | 1730-1888 | 2.93 x10<sup>-8</sup> |
| 15 | Bafer 119 HSP90 | SSA1 66 HSP90 | Unknown | R G M C S T | 405* - 616 | 4.376x10<sup>-06</sup> |
| 16 | AUS1 347 2 HSP90 | UGA 176 HSP90 | SSA1 65 HSP90 | R G M T | 1970 - 2026 | 1.90 x10<sup>-5</sup> |
| 17 | ^NW1 156 2 HSP90 | MED 320 3 HSP90 | AUS1 347 2 HSP90 | G T | 738 - 886 | 1.292x10<sup>-03</sup> |
| 18 | NW1_156_3_HSP90 | Unknown<br>(MEAM1_HSP90_DQ093380) | ASIAII_3_HSP90_HM013713 | _G_ _ _ _ _ | 694 - 852 | 1.352 x10<sup>-3</sup> |

The table represents recombination events, the abbreviations; G- GENECONV, B- Bootscan, M- Maxchi, C- Chimaera and S- Siscan represent recombination detection packages implemented in RDP4.
\* The actual breakpoint position is undetermined (It was most likely over printed by a subsequent recombination event).
^ The recombinant sequence may have been misidentified (one of the identified parents might be the recombinant)
*Unknown* –. The sequence listed as unknown was used to infer the existence of a missing parent sequence. The red colour represents the possibility that an evolutionary process other than recombination could have caused this apparent recombination signal.

**Figure 1.**
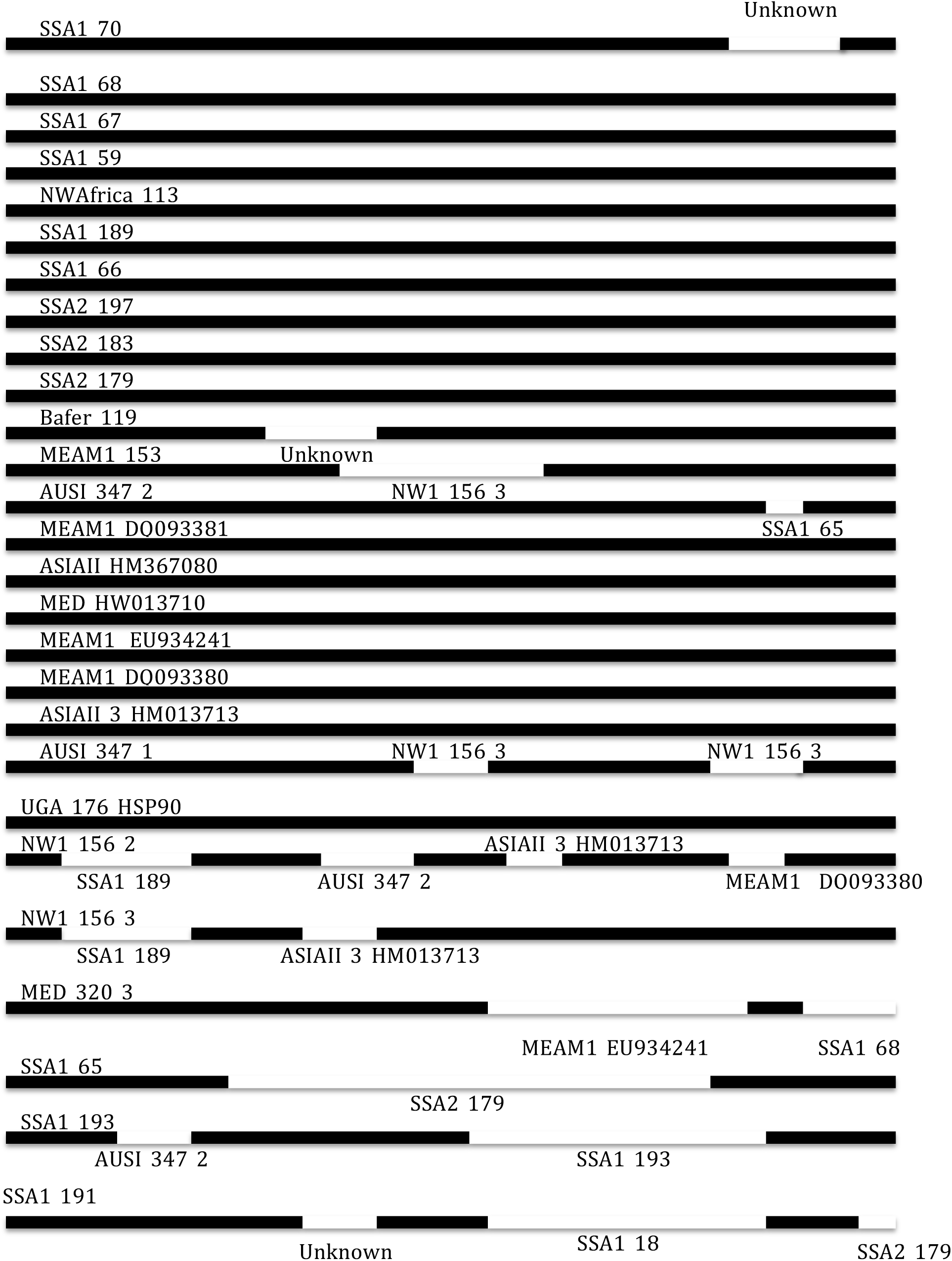
Shows results of the recombination analysis performed on HSP90 sequence alignment of length 2160 base pairs. Recombination events are shown in white and black represents the segment with out recombination events.

Figure 1 shows the results of a recombination analysis of an alignment of 27 sequences of HSP90 gene conducted in RDP4 [36] software. The sequence segments with recombination events are indicated in white and the ones without recombination in black. We observe that most of SSA species did not have recombination events apart from 2 sequences from Uganda (SSA1_70_HSP90 and SSA1_65_HSP90), 2 sequences from Kenya (SSA1_193_HSP90 and SSA1_191_HSP90) and *B. afer* from Malawi. Recombination events were also detected in NewWorld1 (NW1), MED, and AUS1 species. We observed that all the MEAM1 samples previously collected, did not have recombinant sequences apart from the new sample collected from Brazil (MEAM_153_HSP90) and the MED_320_3_HSP90.

Figure 2. Shows the topology of the Bayesian estimate of the phylogenetic tree for the HSP90 gene created from an alignment that is free from recombination events. We observe a long branch leading to clade C that is full of species that had recombination events, this shows that recombination has an effect of the topology of the phylogenetic tree and hence influences evolution. Majority of the species that never had recombination events have short branches as seen in clade A and B. We see sequences from Australia having a relatively long branch in clade B because we detected small segments of recombination in these sequences.

**Figure 2.**
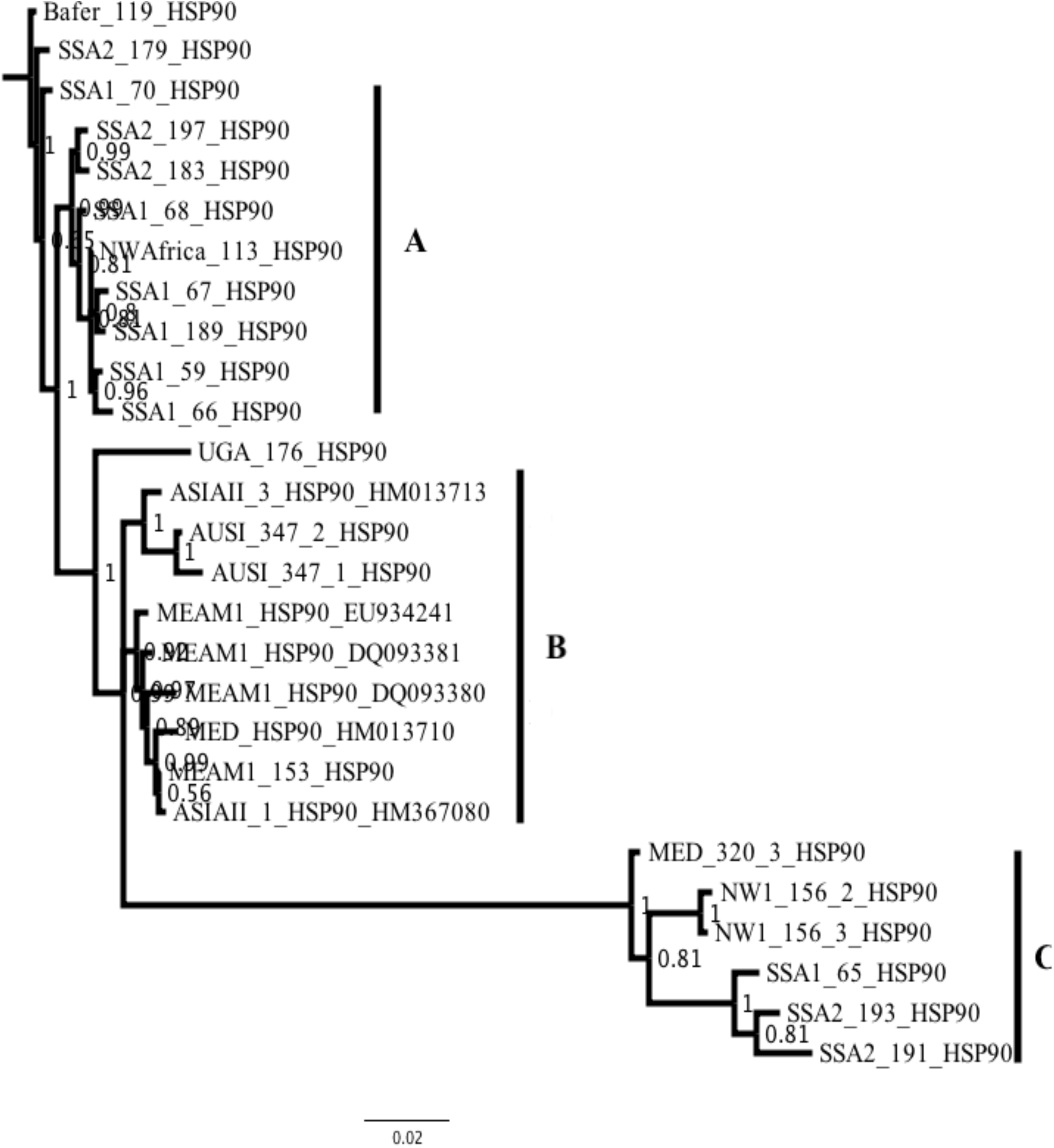
Bayesian phylogenetic tree for HSP90 gene created from an alignment where positions identified as recombinant were removed. A long branch with a mixture of sequences from group A, C and D was formed. The branch length is measured as the expected number of nucleotide substitutions per site.

## DISCUSSION

Recombination events were detected for members of the *B. tabaci* species complex in the HSP90 gene. This could explain the phenomenon of the increased chance of survival and reproductive abundance of whiteflies in hot conditions in East Africa since recombination is a major driving force of evolution [37]. Experimental studies on the members of the *B. tabaci* species complex have demonstrated that HSP plays an important role in thermal adaption, and there is convincing evidence based on HSP90 gene indicating that MED species are adapted to a wider range of temperature than MEAM1 [38-41]. We identified recombinant events in the MED recently collected from the tropical region of Brazil and additional studies show that the cryptic MEAM1 species are being replaced by MED cryptic species in China [39, 42]. Based on our results, we hypothesize the MEAM1 species may not be utilizing recombination in the HSP90 gene to handle climate changes. However, thermal adaptation of whiteflies may depend on many other factors such as, insecticide resistance, host plants, endosymbiont composition and viruses [43, 44]. The invasive MED cryptic species is thought to have originated from North Africa and they could have a connection with the SSA species [45].

We observed that the some SSA species in (Figure 1) had recombination events while others did not have any recombination events, this could be explained by a number of factors, i.e. whitefly host plants, the viruses carried by whiteflies in East Africa, which are different from those in other geographic regions, and lastly, the endosymbiont composition of the whiteflies [46-48]. Further research is needed to evaluate these interesting patterns on the *B. tabaci* SSA species.

Figure 2 represents a Bayesian estimate phylogenetic tree derived from an alignment of sequences without recombination events. We did not estimate the Bayesian phylogenetic tree for the dataset with recombination events because different regions of the alignment might have different underlying evolutionary histories due to recombination [49, 50]. Therefore, ignoring recombination might lead to overestimation of the substitution rate heterogeneity and loss of molecular clock [50]. Usually phylogenetic trees estimated without removing recombination events have larger terminal branches similar to those of exponentially growing populations [50]. This is different for our situation in Figure 2 since, we have short terminal branches. Hence, all portions of recombination events were removed prior to phylogenetic tree estimation. Therefore, recombination drives evolution since it has an effect in the shape of the phylogenetic tree.

Despite the increase in temperature, whitefly numbers have not decreased [39]. We therefore hypothesize that recombination in the HSP90 gene of whiteflies could have played a role in whitefly thermal adaptation. The optimal temperatures for whitefly development ranges from 28.0° C to 32.0° C, which are the current temperatures in the East African region [51, 52]. The chaperon function of HSP90 is also a factor in the survival of whiteflies, as it is triggered by temperature changes [23, 30]. While this is not a conclusive study on climate change in the region, the findings help us understand why there are high numbers of whiteflies in the cassava fields in East Africa, thus forming a basis for future studies. Therefore, we recommend that gene expression studies be carried out on SSA species in East Africa.

## CONCLUSION

Temperature is a key determinant is the geographical distribution and survival of species. Theoretical and experimental results from other studies confirm that HSP90 plays an important role in thermal adaptation but none of these has been conducted on SSA whitefly species. We conclude that, recombination events are present in the HSP90 gene of *B. tabaci* and this could be the major driver for superabundance or adaptive evolution of whiteflies in regions with varying environmental temperatures.

## METHODS

### Whitefly sampling

Seven whitefly samples were collected from whitefly “hot spots” in Uganda in a country-wide survey conducted in 2013 from Nakaseke, Luwero and Nakasongola districts of Uganda. Adult whiteflies from symptomatic cassava were collected from the top five leaves using an aspirator, transferred immediately into 70% ethanol in an Eppendorf tube, and then later exported to the laboratory at the University of Western Australia (UWA) where they were kept at -20° C before analysis. Five whiteflies from Kenya were collected from symptomatic common beans with an aspirator. Collaborators from Brazil, Australia and Malawi also sent whitefly samples already preserved in 70% ethanol and they were stored at – 20° C upon arrival at UWA. See additional file 1.

### Total RNA extraction from individual whiteflies

RNA extractions were performed in the UWA genomics laboratory using an Arcturus PicoPure RNA Isolation Kit (Arcturus, CA, USA) following extraction procedures from [53]. To remove contaminating DNA, DNase and divalent cations from the extracted RNA, we used the TURBO DNA free kit as described by the manufacturer (Ambion Life Technologies CA, USA). To increase the concentration of the RNA, we used a vacuum centrifuge (Eppendorf, Germany) set at room temperature for one hour then resuspended the pellet in 18 µl of RNase free water and stored the prepared RNA immediately at -80° C awaiting further analysis. RNA integrity was quantified by 2100 Bio-analyser (Agilent Technologies).

### cDNA and Illumina library preparation

The Illumina TruSeq Stranded total RNA preparation kit was used to make cDNA libraries from the RNA of each individual whitefly as described by the manufacturer (Illumina, San Diego, CA, USA). Libraries were sent to Macrogen, Korea (www.macrogen.com), where 15 samples were sequenced on a HiSeq 2500 on a rapid run mode and 6 samples on HiSeq 4000 on a high-throughput mode, see additional file 2. Sequencing control software HCS v2.2 and HCS v3.3 were used for base calling and quality assessment respectively.

### NGS data analysis (De novo sequence assembly and mapping)

All samples were trimmed using CLC Genomic workbench 8.5.1 (CLCGW) with the quality score limit set to 0.01, and the ambiguous limit set to 2. Trimmed reads were assembled into contigs using de novo assembly function of CLCGW with automatic word size, automatic bubble size, minimum contig length 500, mismatch cost 2, insertion cost 3, deletion cost 3, length fraction 0.5 and similarity fraction 0.9 (additional file 2). Contigs were imported in Geneious 9.1.8 [54] on Mac OS 10.6, and contigs were then mapped onto the reference sequence for HSP90 MEAM1 EU934241 from GenBank. Mapping was performed with the following settings in Geneious: minimum overlap 10%, minimum overlap identity 80%, allow gaps 10%, fine tuning set to iterate up to 10 times at custom sensitivity. The consensus contig from mapping was aligned to the *de novo* contig using MAFFT [55]. The resulting alignment consensus was manually inspected for ambiguities and gave rise to the new HSP90 sequences covering the full coding region of length 2160 base pairs. Open reading frames (ORFs) were predicted in Geneious and other HSP90 sequences used in this study were downloaded from Genbank (i.e., HM013713, HM367080, HM01370, DQ093380 and DQ093381). Alignment of the new HSP90 sequences with Genbank sequences was performed in Geneious using the MAFFT plugin [55]. A total of 27 sequences of 2160 base pairs were aligned.

### Recombination analysis

An alignment of 27 sequences of the coding region of HSP90 gene was screened for recombination using the recombination detection program RDP4 [36]. An extensive array of methods implemented in RDP4 was used for detecting and visualising recombination events. i.e. RDP [56], GENECONV [57], Bootscan [58], MaxChi [59], Chimaera [60] and SiScan [61]. Default parameter settings were used with a Bonferroni corrected p-value cut off of 0.05. Recombination events detected by four or more methods with significant phylogenetic support were considered reliable evidence for recombination (Table 1). A recombinant free data-set was generated following the recombination analysis by removal of all parts of the alignment that were detected as recombinant events and replaced with gap characters in RDP4 software. The recombination free dataset was later used for phylogenetic analysis.

### Phylogenetic analysis

The recombination free dataset generated from RDP4 software used in phylogenetic analysis. We ran jModelTest 2 [62] on the dataset to select statistically the best-fit model of nucleotide substitution. The best model was GTR+I+G and this model was then used in a Bayesian phylogenetic analysis. We ran MrBayes 3.2.2 [63] in parallel on Magnus (Pawsey supercomputing Centre, Perth, Western Australia) and a phylogenetic trees was constructed. Four Markov chains were run for 50 million generations, trees were sampled every 1000 generations, and 12500 sub-optimal trees were discarded as burn-in at the beginning of the MCMC run. No runs indicated a lack of convergence and the potential scale reduction factor for all parameters approached one. The effective sample size for all parameters was above 200 for each run.

## LIST OF ABBREVIATIONS

ASIAII: Asia II
AUSI: Australia I
CBSD: Cassava brown streak disease
CMD: Cassava mosaic disease
MCMC: Markov chain Monte Carlo
MEAMI: Middle East Asia Minor I
MED: Mediterranean
NWI: New World I
NWAfrica: New World Africa
SSA: Sub-Saharan Africa
UGA: Uganda
UWA: University of Western Australia

## DECLARATIONS

### Ethics approval and consent to participate

Not applicable.

### Consent for publication

Not applicable.

### Availability of data and material

The data that support the findings of this study are available from the corresponding author upon request and sequence data were submitted to Genbank with accession numbers (MH383308 - MH383328).

### Author’s contributions

TK carried out the recombination analysis with inputs from LMB, LSL and AS. TK, BRD, JMW participated in wet laboratory work, TK, JMW, BRD, CAO and TA, provided samples. TK drafted the manuscript and all authors read and approved the manuscript.

#### Acknowledgements

TK is supported by the University of Western Australia, Scholarship for International Research Fees (SIRF), University Postgraduate Awards (UPA) and the University of Western Australia Safety Net top up scholarship. This work forms part of his PhD. The Natural Resources Institute, University of Greenwich, supported sequencing, fieldwork and the first 2 years of TK’s PhD from the grant provided by the Bill and Melinda Gates Foundation (Investment ID OPP1058938). National Crop Resources Research Institute (NaCRRI) provided samples and facilitated sample collection. We thank Paul De Barro, JMW, BRD, and Donald Kachigamba for providing samples. Resources provided by Pawsey Supercomputing Centre with Funding from the Australia Government and the Government of Western Australia supported computation analysis. We thank Joanne Edmondston from Graduate Research School, Joe Bielawski and Laura kubatko for reviewing the earlier versions of this manuscript. Finally, we thank the farmers in East Africa for allowing us to collect samples on their farms.

## TABLE LEGENDS

**Additional file 1.** Table showing species ID, Genbank accession numbers, sample locations and host plants where each species was found

**Additional file 2.** A summary next generation sequencing (NGS) data analysis results in this study

| Sample ID | Number of reads obtained | Reads after trimming | Number of Contigs | Average Contig length | N50 |
| --- | --- | --- | --- | --- | --- |
| 59 | 8,000,000 | 7,848,046 | 16,473 | 1,102 | 1,195 |
| 65 | 8,000,000 | 7,857,579 | 17,146 | 1,024 | 1,071 |
| 66 | 8,000,000 | 7,857,564 | 16,489 | 1,045 | 1,095 |
| 67 | 8,000,000 | 7,876,283 | 18,271 | 1,165 | 1,289 |
| 68 | 8,000,000 | 7,861,938 | 19,852 | 1,201 | 1,368 |
| 70 | 8,000,000 | 7,861,502 | 20,182 | 1,113 | 1,205 |
| 113 | 8,000,000 | 7,850,359 | 13,156 | 1,023 | 1,050 |
| 119 | 8,000,000 | 7,845,846 | 20,368 | 1,238 | 1,400 |
| 153 | 36,449,340 | 36,232,156 | 25,435 | 1,235 | 1,432 |
| 156_2 | 40,826,418 | 40,604,557 | 27,466 | 1,356 | 1,689 |
| 156_3 | 43,194,264 | 42,953,727 | 30,716 | 1,346 | 1,655 |
| 176 | 8,000,000 | 7,835,023 | 19,865 | 1,366 | 1,616 |
| 179 | 8,000,000 | 7,842,509 | 15,570 | 1,194 | 1,295 |
| 189 | 8,000,000 | 7,823,762 | 2,530 | 927 | 910 |
| 191 | 8,000,000 | 7,833,877 | 19,770 | 1,124 | 1,222 |
| 193 | 8,000,000 | 7,852,051 | 20,061 | 1,122 | 1,218 |
| 197 | 8,000,000 | 7,871,950 | 19,224 | 1,230 | 1,408 |
| 320_3 | 39,894,296 | 39,686,374 | 27,867 | 1,144 | 1,250 |
| 347_1 | 39,142,358 | 38,894,512 | 30,496 | 1,372 | 1,723 |
| 347_2 | 37,306,246 | 37,071,584 | 27,343 | 1,314 | 1,614 |
| 183 | 8,000,000 | 7,816,022 | 20,066 | 1,298 | 1,512 |

| Sample ID | Species ID | Genbank Accession # | Country | Latitude | Longitude | Altitude | Host plant |
| --- | --- | --- | --- | --- | --- | --- | --- |
| 59 | SSA1 | MH383311 | Uganda | 0.85070 | 32.41920 | 1078 | Cassava |
| 65 | SSA1 | MH383326 | Uganda | 0.87345 | 32.64796 | 1085 | Cassava |
| 66 | SSA1 | MH383314 | Uganda | 0.86932 | 32.51936 | 1113 | Cassava |
| 67 | SSA1 | MH383310 | Uganda | 1.41966 | 32.43936 | 1064 | Cassava |
| 68 | SSA1 | MH383309 | Uganda | 1.19876 | 32.73804 | 1069 | Cassava |
| 70 | SSA1 | MH383308 | Uganda | 1.31247 | 32.58484 | 1042 | Cassava |
| 113 | NWAfrica | MH383312 | Uganda | 0.54896 | 31.44420 | 1283 | Mujjaja |
| 119 | *B. afer* | MH383318 | Malawi | -12.792975 | 34.200964 | 500 | Cassava |
| 153 | MEAM1 | MH383320 | Brazil | -22.844553 | -48.435439 | 800 | Cabbage |
| 156_2 | NWI | MH383323 | Brazil | -21.980278 | -50.788312 | 397 | Euphorbia heterophylla |
| 156_3 | NWI | MH383324 | Brazil | -21.980278 | -50.788312 | 397 | Euphorbia heterophylla |
| 176 | UGA | MH383319 | Kenya | 00.00622 | 034.73450 | 1510 | Common bean |
| 179 | SSA2 | MH383317 | Kenya | 00.02544 | 034.62852 | 1561 | Common bean |
| 189 | SSA1 | MH383313 | Kenya | 00.33262 | 034.25285 | 1322 | Common bean |
| 191 | SSA2 | MH383328 | Kenya | 00.56222 | 034.17064 | 1165 | Common bean |
| 193 | SSA2 | MH383327 | Kenya | 00.61071 | 037.37062 | 1167 | Common bean |
| 197 | SSA2 | MH383315 | Kenya | 00.59737 | 034.43888 | 1331 | Common bean |
| 320_3 | MED | MH383325 | Brazil | -23.01694 | -048.06861 |  | Cassava |
| 347_1 | AUSI | MH383322 | Australia |  |  |  |  |
| 347_2 | AUSI | MH383321 | Australia |  |  |  |  |
| 183 | SSA2 | MH383316 | Kenya | 00.00622 | 034.55100 | 1561 | Common bean |

